# Exploring the Brain Characteristics of Structure-informed Functional Connectivity through Graph Attention Network

**DOI:** 10.1101/2023.11.30.569343

**Authors:** Zifan Wang, Paule-J Toussaint, Alan C Evans, Xi Jiang

**Author notes:** Corresponding author at: Montreal Neurological Institute, McGill University, Montreal, QC, Canada, H3A 2B4 and School of Life Science and Technology, University of Electronic Science and Technology of China, Chengdu, China.

## Abstract

Independent brain regions in neuroanatomy achieve a specific function through connections. As one of the significant morphological features of the cerebral cortex, previous studies have found significant differences in the structure and function of the cerebral gyri and sulci, which provides a basis for us to study the functional connectivity differences between these two anatomic parts. Previous studies using fully connected functional connectivity (FC) and structural connectivity (SC) matrices found significant differences in the perspective of region or connection in gyri and sulci. However, a clear issue is that previous studies have only analyzed the differences through either FC or SC, without effectively integrating both. Meanwhile, another nonnegligible issue is that the subcortical areas, involved in various tasks, have not been systematically explored with cortical regions. Due to the strong coupling between FC and SC, we use SC-informed FC to systematically explore the functional characteristics of gyri/sulci and subcortical regions by combining deep learning method with magnetic resonance imaging (MRI) technology. Specifically, we use graph attention network (GAT) to explore the important connections in the SC-informed FC through the Human Connectome Project (HCP) dataset. With high classification results of above 99%, we have successfully discovered important connections under different tasks. We have successfully explored the importance of different types of connections. In low threshold, gyri-gyri are the most important connections. With the threshold increasing, sub-sub become the most important. Gyri have a higher importance in functional connectivity than sulci. In the seven task states, these connections are mainly distributed among the front, subcortical, and occipital. This study provides a novel way to explore the characteristics of functional connectivity at the whole brain scale.

## Introduction

Although the functions associated with specific brain regions have been well-established (Penfield & Boldrey, 1937; Van Essen, 1979; Keller, Crow, Foundas, Amunts, & Roberts, 2009; Amunts & Zilles, 2015; Binder, 2015), it is important to note that no brain region works in isolation, and specific tasks cannot be accomplished solely by individual regions. The brain functions as a complex system that relies on the coordinated transmission of messages between interconnected brain regions, collectively impacting and determining brain function (K. Friston, 2002; Cole & Schneider, 2007; K. J. Friston, 2009; Duncan, 2010; K. J. Friston, 2011; Du et al., 2020). For instance, Pascale Tremblay has emphasized the difficulty that classical language models face in integrating white matter connectivity with the language system. Language networks are inherently distributed structures, and classical models have often overlooked the connectivity between the cerebral cortex and subcortical regions. However, pathways linking language processes to subcortical areas may play a more prominent role (P. Tremblay & Dick, 2016). João D. Semedo’s research has shown how visual signals are transmitted through activity patterns of feedforward and feedback interactions between V1 and V2 (Semedo et al., 2022). Neuroscientists are now considering a new perspective to explore the brain function known as the connectome (Sporns, Tononi, & Kötter, 2005; Thiebaut de Schotten & Forkel, 2022). The connectome examines brain connections at multiple scales, from microscopic levels such as molecular and cellular to macroscopic levels such as brain regions. The macro connectome can effectively reveal the patterns of connectivity between different brain regions, allowing for investigating the functional characteristics of the brain during various tasks (Axer & Amunts, 2022).

The development of connectome cannot be separated from the progress of magnetic resonance imaging (MRI). Through diffusion MRI (dMRI), it is possible to reconstruct the neural pathways of the brain and use methods such as graph theory to study the properties of the brain connectome (Bullmore & Bassett, 2011). Meanwhile, functional MRI (fMRI) are increasingly utilized to explore the functional characteristics of the brain non-invasively. Following Bharat Biswal’s discovery of high correlation in spontaneous coherence in the motor cortex between the left and right hemispheres during resting state Fmri (rs-fMRI) (Biswal, Zerrin Yetkin, Haughton, & Hyde, 1995), numerous studies have confirmed the ability of fMRI to identify functional differences between various brain regions (Barch et al., 2013) as well as the existence of brain networks where regions exhibit potential functional relevance in a resting state (Damoiseaux et al., 2006; Yeo et al., 2011). For example, Thomas Yeo divides the brain into seven or seventeen different functional activity areas through rs-fMRI, which are widely distributed anatomically in the brain. Task-based fMRI (ts-fMRI) enables the examination of activation patterns in specific brain regions during specific tasks, providing valuable insights into the neuroimaging underlying brain function at the regional level (K. J. Friston, 2009). However, the completion of brain function is inseparable from the transmission of information through structures. The nerve fibers inside the brain are responsible for the transmission of electrical signals between brain regions, thereby mediating various functions. There are many methods to study the relationship between function and structure, and one of the most extensive methods is to generate the FC through fMRI and SC through dMRI of the brain. It is evident from various studies that there is an high coupling between the structure and function of the brain and the value is significant positive (Damoiseaux & Greicius, 2009; Suárez, Markello, Betzel, & Misic, 2020). Thus, it is crucial to integrate both structure and function to fully explore the characteristics of the brain.

The generation of FC and SC is based on various brain atlases. In recent studies, researchers have proposed several hypotheses regarding the formation of the highly folded topology of the brain surface, characterized by gyri and sulci. These hypotheses include external factors such as the constraints imposed by the brain skull (Welker, 1990; Miska et al., 2004), internal factors such as differential laminar growth (Richman, Stewart, Hutchinson, & Caviness Jr, 1975), genetic regulation (Mota & Herculano-Houzel, 2012), axonal tension or pulling (Essen, 1997; Nie et al., 2012). Although the exact mechanism behind cortical folding remains unknown, notable anatomical (Fischl & Dale, 2000; Smart, Dehay, Giroud, Berland, & Kennedy, 2002) and morphological (Magnotta et al., 1999) differences have been identified between gyri and sulci. Building upon these foundations, researchers have divided brain regions into gyri and sulci in various brain atlases, revealing functional (Liu et al., 2019; X. Jiang, Zhang, Zhang, Kendrick, & Liu, 2021; M. Jiang et al., 2022), and structural (Deng et al., 2014) distinctions. For example, Deng found significant differences in the strength of functional connections between gyri and sulci, with gyri-gyri connections exhibiting the strongest functional connectivity and sulci-sulci connections displaying the weakest (Deng et al., 2014). Jiang observed an abundance of heterologous functional regions and spatial overlap patterns of functional networks in the gyri compared to the sulci (X. Jiang et al., 2015; X. Jiang et al., 2016). Recently, it has been discovered that the important functional connectivity between gyri is greater than that between sulci during different task states (M. Jiang et al., 2022). However, these studies primarily focus on exploring the functional characteristics of the brain through gyri and sulci, neglecting the potential role of subcortical regions in various tasks. Additionally, these studies mainly employ fMRI data to investigate brain function, without effectively integrating the structural information of the brain from dMRI with fMRI data to examine functional characteristics under different task states. Given the stability of the brain’s structural connections, how do these invariant structural connections give rise to diverse functions? In this study, we aim to address this question by using the structural connectivity matrix to sparsity the fully connected functional connectivity matrix. Our goal is to explore the similarities and differences in connectivity patterns between structurally connected brain regions during different tasks.

Due to the non-topological structure of the brain, we use a deep learning model based on graph attention network and use the functional connection matrix constrained by the structural connection matrix as the input to explore the functional brain network in each task state compared with the resting state. Our model can accurately distinguish the resting state from the task state with an accuracy of around 99%. Then we use the interpretability algorithm based on gradient to extract the important functional connections in each region, which constitute the most important brain network in the task state. We found that under the different thresholds of the seven task states, for cortical connections, the gyri-gyral connections are the most important, gyri-sulcal connections in the middle and sulci-sulcal connection at the least. While from another perspective of whole brain scale connection, the gyri-gyral connections are the most important at the very beginning then decreases and sub-sub connections increase their importance with the thresholds increase and reaches peak around the top 5 percent. At the same time, we found that gyri-sub connections are also more important than sulci-sub connections. This shows the importance of gyri in the task state and the extensiveness of the subcortical areas. By computing the degree of each brain region, we found that frontal lobe is the most important lobe in the brain.

Overall, our work has the following contributions:

1. Inspired by the high coupling relationship between tasks and structures, we use SC to sparse the FC matrix and explore important structural connections under each task.
2. We systematically explored the functional connectivity patterns of the brain from the perspective of the whole brain, including gyri, sulci, and sub.
3. Due to the non-Euclidean properties of the brain, we use graph attention networks to explore the connectivity characteristics between gyri, sulci, and sub.

## Materials and Methods

### 2.1 Participants

We use Human Connectome Project Young Dataset (HCP-YA) (Van Essen et al., 2012; Matthew F. Glasser et al., 2013; Van Essen et al., 2013; Elam et al., 2021) in our study. The HCP-YA dataset include 1206 subjects originally, after selecting subjects who has data for all one resting state and seven task states (Barch et al., 2013), and a complete structural connectivity (SC) matrix, a total 999 subjects are include here, aging from 22 to 37 years (mean ± SD age of 28.71 ± 3.69 years). All subjects are scanned using a customized Siemens 3T “Connectome Skyra” with a standard 32-channel Siemens receive head coil at Washington University in St. Louis. The detailed parameters about the T1w, T2w, resting state fMRI, task state fMRI, and diffusion imaging can be find here (Wu-Minn, 2017).

The resting state has a run duration of 14:33 minutes and number of frames of 1200, and four runs for each subject. Here we only use the REST1 of RL. The seven tasks (working memory, gambling, motor, language, social, relational, and emotion) have different run durations and number of frames: 5:01 minutes and 405 frames, 3:12 minutes and 253 frames, 3:34 minutes and 284 frames, 3:57 minutes and 316 frames, 3:27 minutes and 274 frames, 2:56 minutes and 232 frames, and 2:16 minutes and 176 frames, respectively.

### 2.2 Data preprocessing

To conduct functional connectivity (FC) matrix, first we extract signals from the datasets. We utilize ICA-FIX MSMAll grayordinate data, which has already register signals from voxel to cortical brain, making an easier and efficient way to analysis brain function (Marcus et al., 2013). The HCP dataset has already done minimal preprocessing and can be directly downloaded from the HCP website at https://www.human-connectome.org/. The minimally preprocessing procedure involve the following steps: prefreesurfer, freesurfer, postfreesurfer, fmrivolume, fmrisurface, ica-fix, and msmall, more detailed information can be found in this paper (Matthew F Glasser et al., 2013).

After we extracte brain signals from the from the grayordinate data, which consists of 96854 vertex and pixel measurements. Specifically, 64984 vertexes are corresponding to cortical regions and 31870 voxels are corresponding to subcortical regions. We use Desikan-Killiany (DK) atlas to define the brain into regions of interest (ROI) (Desikan et al., 2006). Since DK atlas has only cortical regions parcellation, we use the automatic parcellation atlas provided by Freesurfer which consist of 19 regions for subcortical regions. We then divide the cortical brain regions into gyral/sulcal areas based on the sulc value (Fischl, Sereno, & Dale, 1999). For cortical regions, positive sulc values represent gyri, while negative values represent sulci. Gyri correspond to the convex parts of the brain surface, while sulci represent the concave parts. Based on the previous study, the selected top or bottom value has no influence on the consistency of the final result (Yang et al., 2019). Here, we select the top and bottom 20% of sulc values in each region to consider them as gyri or sulci. Unfortunately, subcortical regions cannot be divided into gyral/sulcal units due to technical shortcomings. By applying the thresholds, we successfully determine whether a region corresponds to a gyral or sulcal area, which made us get 155 ROI for the whole brain scale (Fig. 1A).

**Fig. 1.**
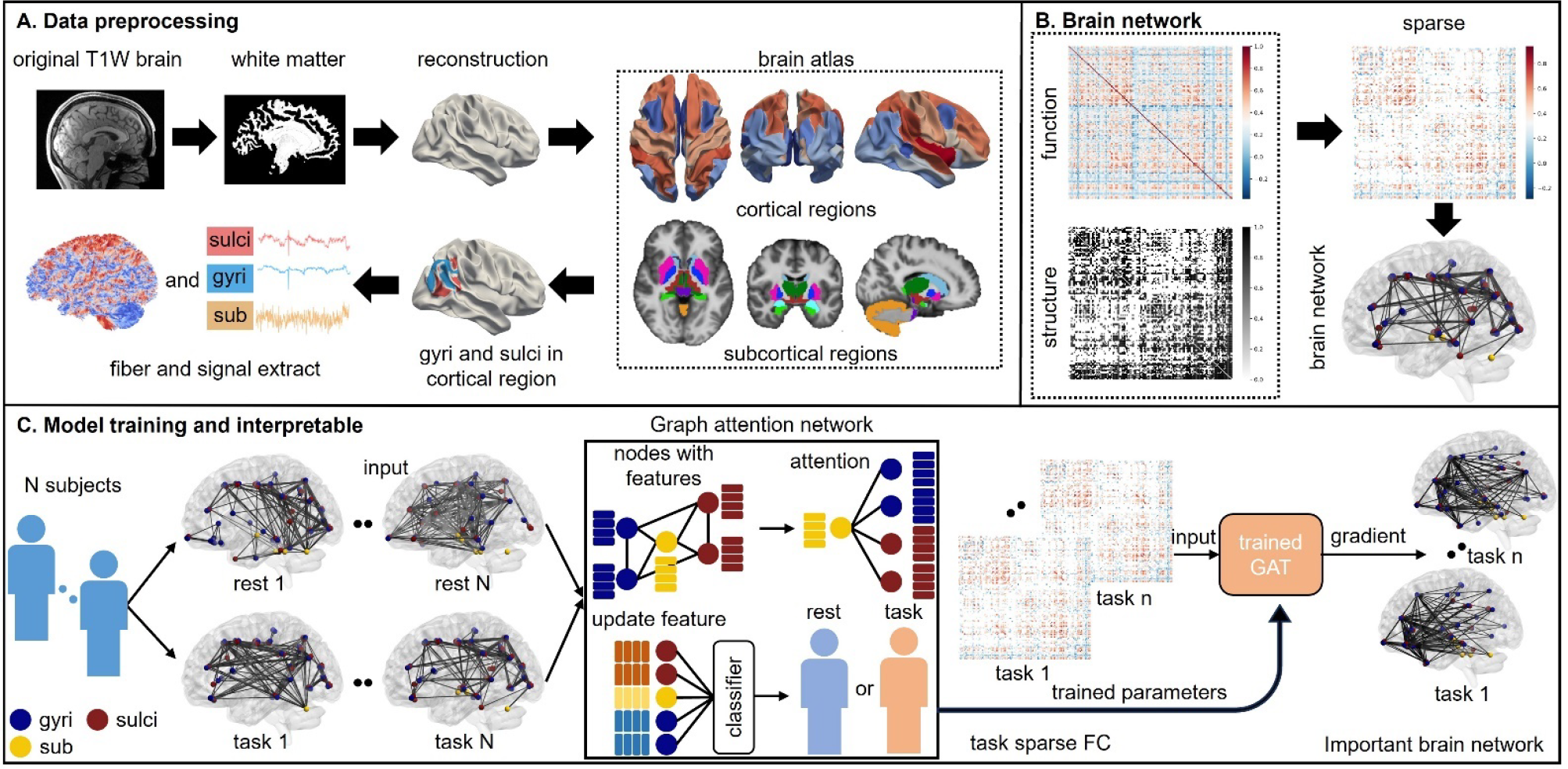
Overview of our method. Fig. 1A shows the preprocessing process of our dataset. We extract brain function and structure data from fMRI data and dMRI data, respectively. Fig. 1B shows the construction of our input data. Using structural connectivity as an adjacency matrix to sparse the fully connected functional connectivity, we construct the brain network for each subject in each state. Fig. 1C shows the overview of our model and the interpretable method. We use graph attention network to do a binary classification and by using a gradient based method, we can explore the gradient of each connection and consider it as the importance.

To compute the functional similarity between brain regions, we calculate the average signal within each ROI. Subsequently, we employ the widely used Pearson correlation to quantify the functional connectivity between two ROIs by using their average signal:

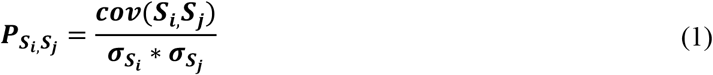

Where the *S*_*i*,_*S*_*j*_ ∈ *S* = *S*_1,_*S*_2_, … *S*_*n*_ (*n* = *num of cortical regions*) represents individual brain region and *cov*(.) and *σ*_(.)_ represent the covariance and standard deviation respectively. For each subject, we could calculate their one rest and seven tasks functional connectivity matrix (rs-FC and ts-FC).

To generate the structural connectivity (SC) matrix, we follow most of the procedures outlined in the BATMAN pipeline (Tahedl, 2020). However, since our objective is to construct a SC matrix with division based on gyri and sulci, we need to first create an individual annotation file which is parceled into gyri and sulci. The annotation file is initially in the fsaverage space, so we convert it into the fs_32k space. Next, we use the sulc value to partition the cortical surface into gyri and sulci. Finally, we reconvert the fs_32k space back to the fsaverage space to obtain the desired annotation file. It is important to note that there may be slight variations in the cortical surface, including parcellation and sulc values, among different subjects. Therefore, each subject requires an individualized annotation file.

Once we acquire the individual annotation files for each subject, we follow the procedures outlined in BATMAN to successfully construct the SC for each subject. We then utilize the SC matrix to sparsity both the resting state functional connectivity (rs-FC) and task state functional connectivity (ts-FC) of each subject.

Specifically, we set the connections with non-zero structural connection values to 1 to create a sparse SC matrix. This sparsity mask is subsequently applied to the functional connectivity data, resulting in sparse functional connectivity (sparse FC) (Fig. 1B).

### 2.3 Graph Attention network and its interpretability

Due to the unique non-Euclidean geometric structure of the brain and the fact that current modeling and analysis of the brain mainly rely on ROI selection and connections between ROIs, convolutional neural networks (CNN) with high accuracy in identifying Euclidean structured images cannot be well applied to such brain data (He, Zhang, Ren, & Sun).

Graph attention network (GAT) is a specialized network for processing graph data (Velickovic et al., 2017), and its unique attention mechanism makes it more flexible in application compared to graph convolutional network (GCN). The traditional GCN is a type of neural network that processes graph structured data in the frequency domain, but it requires the structure of each input graph to remain invariant (Kipf & Welling, 2016; S. Zhang, Tong, Xu, & Maciejewski, 2019). In our experiment, as the SC matrix of each individual may be different, leading the sparse FC matrix may also have different connection relationships. At the same time, due to the attention mechanism being able to act on all nodes that are related to the ROI and calculate the attention score between the current ROI and other nodes, this mechanism is very suitable for exploring the importance of edges connected between nodes. Specifically, since each subject has a resting state and seven task states, we could obtain the structural brain network based functional connectivity values (S-FC) of each participant in eight states. Next, we input the S-FC matrix of all individuals in their resting state and one task state into GAT each time and perform a 5-fold cross validation under six different random seeds. Finally, we obtained 30 trained model parameters for each task state.

Our GAT is composed of two stacked graph attention layers (GAL), each divided into eight attention heads. Specifically, this is how we implement our graph attention network. First, for each subject, the node feature of each ROI is based on the S-FC, thus each node has 155*1 node features here. Then for each ROI, it only calculates the attention coefficient between nodes connected to it. Specifically, the formula for attention calculation is as follows:

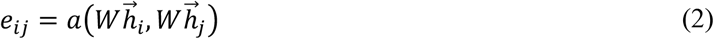

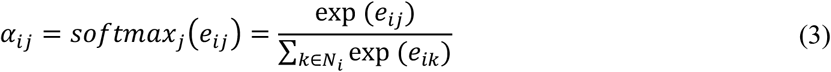

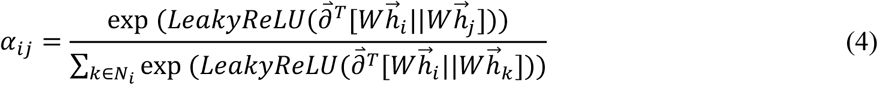

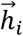 and 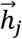 is the node feature of*node*_*i*_ and 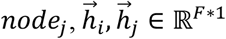. *W* ∈ ℝ^*F*′∗*F*^ is a trainable parameter that maps the features of each node from *F* to *F*^′^, thus 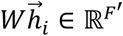. *a* represents attention mechanism between *node*_*i*_ and *node*_*j*_ and *e*_*ij*_ is the score of the importance of *node*_*i*_ and *node*_*j*_. *α*_*ij*_ represent the attention value after SoftMax. Since each node may be connected to multiple other nodes at the same time, the SoftMax function is used here to normalize the attention coefficients of *node*_*i*_ with other nodes to between 0 and 1, while adding them to 1. Specifically, for the calculation of attention coefficients between two nodes, we concatenate the features of each node after *W* projection to form a 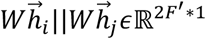 feature. Using 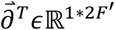 to do matrix multiplication with features after concatenating, we obtain a *scoreϵ*ℝ^1∗1^. Finally, by using the LeakyReLU, we obtained the attention coefficient between the two nodes.

Another major advantage of the GAT model is the use of a multi head attention mechanism to ensure the stability of the attention obtained by the model. Specifically, we use eight attention heads, each of which is independent, so we can obtain eight different attention scores, add them up or average them, and ultimately obtain our results. Therefore, in each layer of GAL, the features of each node are assigned to eight attention heads. In the first layer of GAL, each attention head projects features from 155 to 384, while in the second layer of GAL, the feature dimensions remain unchanged. Finally, by adding a classification layer, we obtain the classification results of the model.

After training the model, we use a gradient based approach to determine the important connections that the model focuses on (Chen et al., 2022). Specifically, we input the brain network features and actual labels of each individual under a certain task, and by calculating the gradient of predicted values under each feature, we can obtain the importance of each feature relative to that category. Since our feature input is a S-FC between brain regions, we can obtain the importance of each connection of S-FC in that category. By adding the results of all individuals, we obtain a group average result as our final output. The specific formula for exploring the weight of edges relative to a certain category is as follows:

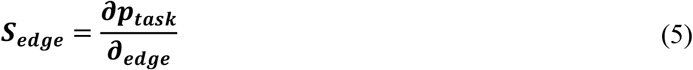

Where *S*_*edge*_ is the gradient value of each connection for that category. We then can choose top k connections to conduct a functional brain network.

### 2.4 Adversarial Training in GAT

By adding the calculated perturbation noise to the original image, the classifier may can correctly classify the original image and incorrectly classify the image with the perturbation even the amplitude of this perturbation is very small (Goodfellow, Shlens, & Szegedy, 2014; Miyato, Dai, & Goodfellow, 2016). Human eye will not figure out these pictures, but it can easily “trick” the deep neural network during the classification stage. Hence, we add adversarial training in our model to increase the robustness of our model. In addition, some previous work indicates that gradient in the model with adversarial training is more stable and the visualization result is also more stable (T. Zhang & Zhu). Thus, adding adversarial training can benefit the interpretability of model. Here we use Fast Gradient Method (FGM) (Goodfellow et al., 2014) to generate adversarial samples. The constrained perturbation can be defined as:

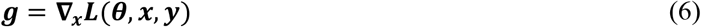

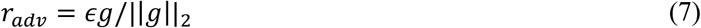

Where the *θ* is parameters of a model, x is the input of a model and y is the target associated with x. And the *L*(*θ, x, y*) is the loss of the model and ∇_*x*_ represent the derivation of x for *L*(*θ, x, y*). The basic idea of adding perturbation is following the direction of the gradient ascent, and for every step, we used the *L*_2_ norm to constrain every step to get a unique *r*_*adv*_ for every training. We (1) calculated the forward loss of x and based on back propagation to obtain a gradient. (2) Then *r*_*adv*_ was calculated from the gradient of the embedding layer and add to the current embedding. (3) Then we got the adversarial sample *x* + *r*_*adv*_ and calculated the forward loss of *x* + *r*_*adv*_, and the back propagation got the gradient which will add it to the original gradient of x. (4) Then we restored the parameters of embedding dim at the original value. (5) Finally, the parameters were updated according to the gradient of (3). We set the perturbation factor as 0.001 here.

### 2.5 Analysis of important connections

After obtaining the connection weights, we conduct analyses to assess the stability and repeatability of our findings using different thresholds and model parameters. Initially, we compute the average results across thirty model parameters by aggregating the connection values obtained under each parameter. We employ various thresholds to determine the number of connections for quantitative analysis. There are six types of connections: gyri-sulci, sulci-sulci, gyri-gyri, sulci-sub, gyri-sub, and sub-sub. Considering the varying number of connections for each type of connection, we calculate the number of connections for each type among all individuals and added them together to determine the normalization coefficient. Specifically, the normalized data would be as follow:

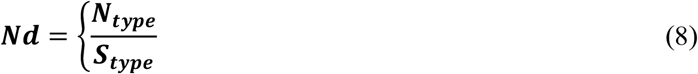

Where *Nd* is the normalized data, *N*_*type*_ represents the number of a certain type of connection and *S*_*type*_ represents the real number of this type in SC.

Upon obtaining these crucial connections, we further analyze their distribution within the brain regions, which can be categorized into six lobes (frontal, limbic, temporal, parietal, occipital and subcortical regions). By leveraging the concept of node degree from graph theory, we determined the importance of these regions. We can also explore the importance of gyri, sulci and subcortical regions following the same procedure. Finally, we analyze the most important brain regions under different thresholds.

### 2.6 Model Training Scheme and Parameter Setting

We adopt 6 times 5-fold cross-validation strategy to ensure the robustness of the results. All subjects are split into 800 as training sets and199 as test sets in each fold respectively. We use the CrossEntropy as the loss function and Adam as the optimizer. We set the learning rate to 0.001 with a decay rate weight of 0.0001, the dropout rate as 0.5 to reduce the overfitting, the batch size to 32, the total quantity of epochs to 60, the early stopping epochs to 7. The model training is based on the Pytorch framework with a GTX3090 GPU.

## Result

### 3.1 Similarity and differences of S-FC among seven tasks

Firstly, we perform five-fold cross-validation under six random seeds and calculate the average results. We examine different thresholds ranging from top 0.1% to top 1% with an interval of 0.1%, and from top 1% to top 20% with an interval of 1% to validate our conclusions. After dividing the connections into six types and standardizing them, we present our results in Figure 2.

**Fig. 2.**
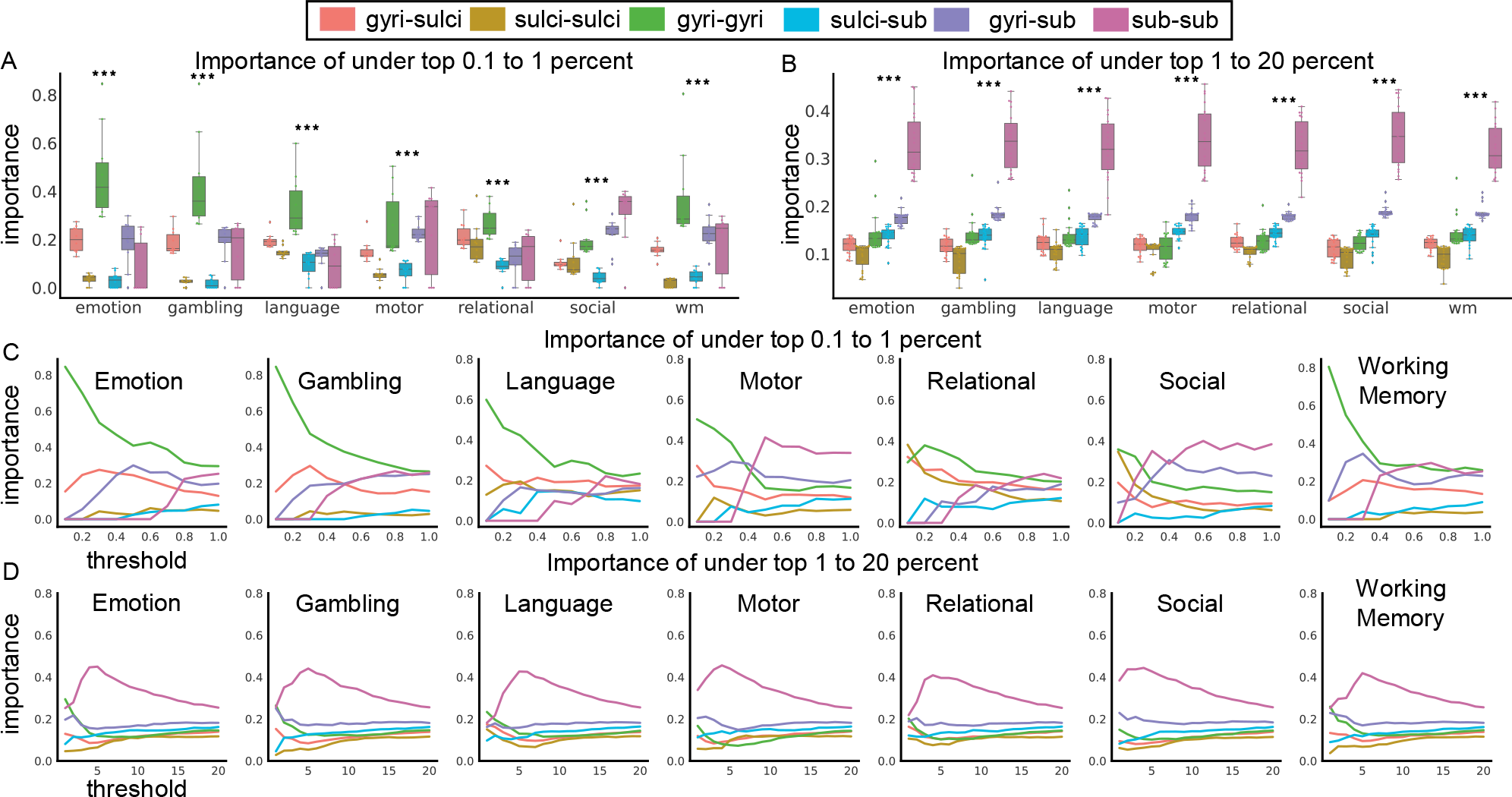
Stability of our conclusion under different thresholds. We use two different thresholds to verify our conclusion, a low threshold and another relatively high threshold. Fig. 2A, B show the importance of the six connections within the current threshold range. Fig. 2C, D show the line plot under each threshold. We do a one-way ANOVA in each task state, and *** indicates p < 0.001.

We observe that when the threshold range is relatively low, gyri-gyral connections play the most important role in the seven task states, except in the motor task (Fig. 2A). However, in the motor task, the most important connections at the beginning are gyri-gyri (Fig. 2C). Additionally, while focusing solely on cortical connections, gyri-gyral connections are found to be the most important, followed by gyri-sulcal connections and then sulci-sulcal connections. Moving on to cortical-subcortical connections, we observe that gyri-sub connections hold greater significance than sulci-sub across all seven tasks. Additionally, the importance of gyri-gyral connections decrease as the threshold increase (Fig. 2C). When examining higher thresholds, we find that although the relationship among cortical-cortical and cortical-subcortical connections remained unchanged, the gyri-gyral connections are no longer the most crucial connections in the entire brain. Instead, sub-sub connections emerge as the most important during the seven task states (Fig. 2B). Meanwhile, the connections between cortical-subcortical and cortical-cortical are the three most important types of connections. As the threshold increased, the importance of sub-sub connections peaks around the top 5%, after which it gradually decreases. Notably, the sub-sub connections consistently maintain as the most important connection across different threshold ranges (Fig. 2D).

Next, we visualize the important connections under different thresholds and divided the cortical regions into five different lobes: frontal, limbic, temporal, parietal, and occipital. Taking emotion as an example to verify the stability of important connections under different thresholds, we find that as the threshold increases, the distribution pattern of important connections in the brain is stable, and the main areas of connections are frontal, limbic, and subcortical regions (Fig. 3A).

**Fig. 3.**
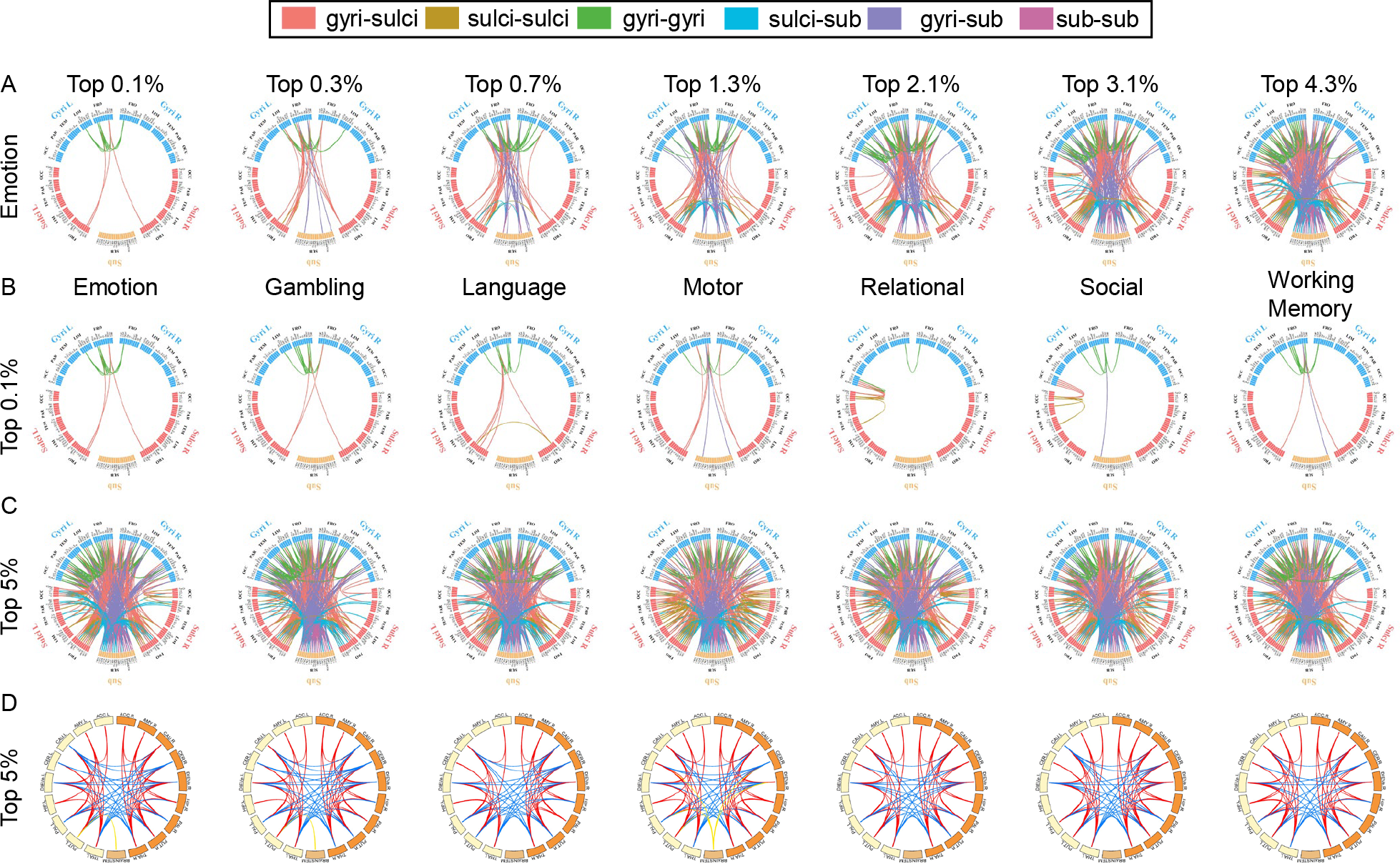
Visualization results of important connections under different thresholds. Fig. 3A shows the connections under seven different thresholds in emotion task. Fig. 3B shows the top 0.1% percent connections in seven tasks. Fig. 3C shows the top 5% connections in seven tasks. Fig. 3D shows the sub-sub connections in seven tasks, red lines represent intra-hemispheric connections and blue lines represent inter-hemispheric connections, yellow lines represent connections with brainstem.

We show two different results from top 0.1% (gyri-gyral occupies the most important position), and top 5% (sub-sub occupies the most important position). In top 0.1% connections, we find that frontal, limbic, and subcortical regions are the main regions where these connections locate in emotion, gambling, language, motor and working memory. While in social and relational, connections between occipital regions are also important. In top 5% connections, we find that almost all brain regions are involved, and the distribution of connections in the seven task states are relatively similar. Due to the highest importance of the sub-sub connections, we have shown the distribution patterns of sub-sub connections in Fig. 3D. Under seven tasks, the connection patterns in the intra-hemispheric tend to be symmetrical and are similar among seven tasks. At the same time, for inter-hemispheric connections, there are fewer connections between areas responsible for the same function, and this connection tends to be connected to different functional areas.

Next, we want to explore the common connections among the seven task states under different thresholds. As the thresholds increase, the ratio of common connections among seven tasks increases (Fig. 4A). Under low threshold condition, the common ratio is relatively low. However, when selecting the top 3000 (around top 20%) important connections, the ratio in the seven task states is close to 1, revealing the uniqueness and commonality of connections in the seven task states. Under different tasks, the most important connection patterns corresponding to the task are different, but compared to the resting state, the brain has consistency in important connections when completing the task. Next, we visualize these common connections under different thresholds. The detailed visualization connection pattern is selected at top 0.5 percent. At this point, there are 15 shared connections, accounting for 25% of the total number of connections (Fig. 4B). We find that the common connection types belong to gyri-gyri, gyri-sulci, and gyri-sub, which not only reveals the importance of gyri cross task states, but also its connection patterns indicate that gyri are more involved in advanced functional cognition. At the same time, the left brain is more involved in connections in the seven tasks, which may be related to brain lateralization (Nielsen, Zielinski, Ferguson, Lainhart, & Anderson, 2013). Similarly, we found that the distribution patterns of common connections exhibit stability under different thresholds (Fig. 4C), with the main type being gyri-gyri, gyri-sulci, and gyri-sub. Meanwhile, these connections are mainly located among frontal, limbic and subcortical regions.

**Fig. 4.**
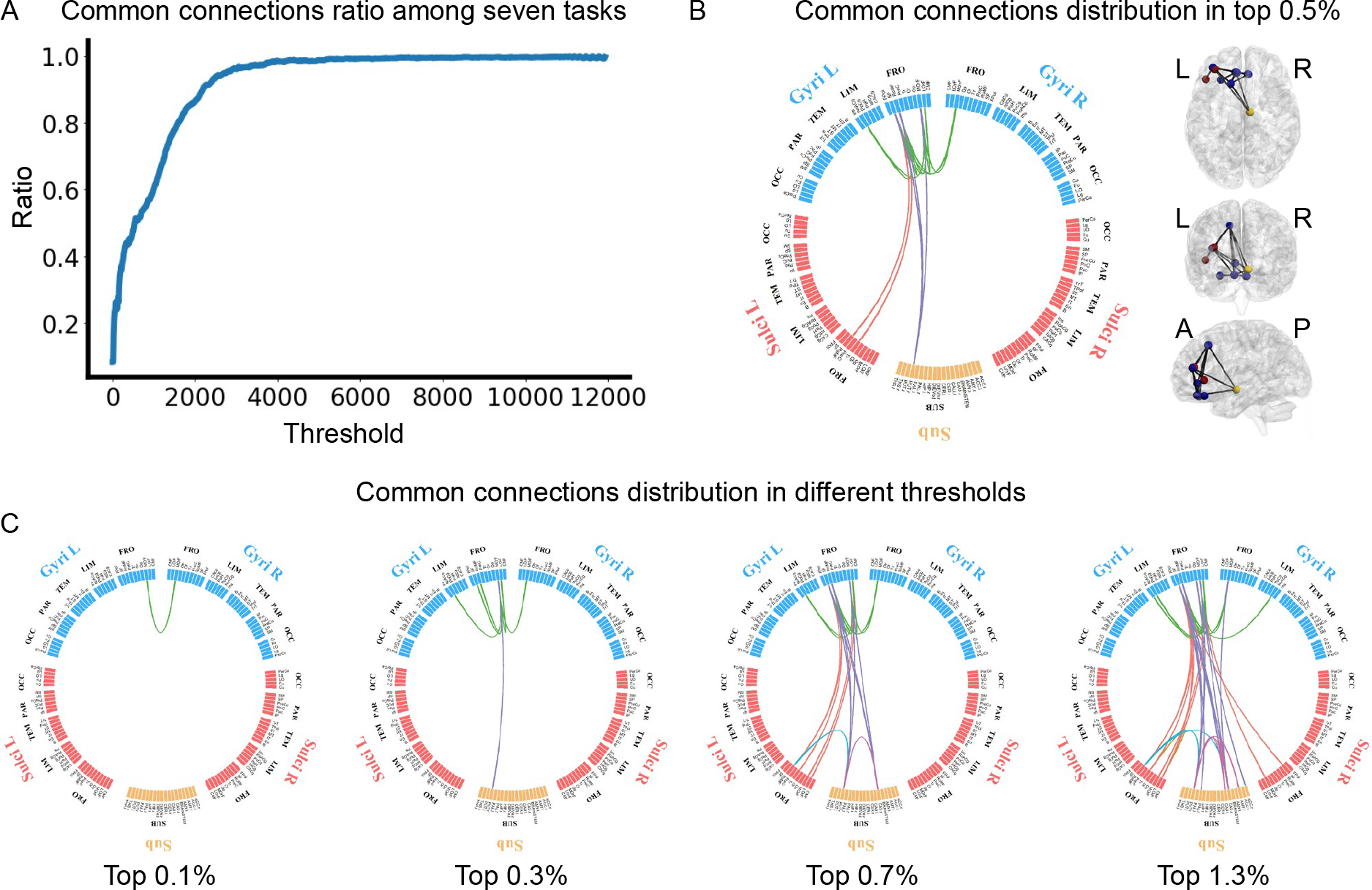
Common connections among seven tasks. Fig. 4A shows the ratio of common connections among seven tasks. Fig. 4B shows the distribution pattern of common connections at thresholds of 0.5%. Fig. 4C shows the stability of common connections’ distribution pattern under different thresholds.

### 3.2 Important brain regions in connectivity

We conduct a quantitative analysis, using degree in graph theory to measure the importance in connectivity. First, we explore the distribution of connections in six lobes (five cortical lobes and subcortical lobe) and mapping these connections into these lobes to compute the degree of each lobe with different thresholds. The degree in each lobe is standardized then. At low thresholds, frontal, limbic, and subcortical lobes are the most important lobes among emotion, gambling, language, motor and working memory, where there is a difference in relational and social, occipital frontal and subcortical are the most important. With the threshold increase, subcortical lobe become the most important while the importance of parietal gradually increases. Finally, the importance of the six lobes trend to be stable in seven tasks, the order from largest to smallest is subcortical, parietal, frontal, occipital, limbic and temporal (Fig. 5A). When considering brain regions into gyri, sulci, and sub, we find gyri have the most importance when the threshold is low, and it gradually decreases with the threshold increasing. While sulci have the least importance in most all thresholds and subcortical regions have less importance when the threshold is low compared to gyri, but it gradually reaches its peak around top 5 percent and become the most important regions among the three parts (Fig. 5B).

**Fig. 5.**
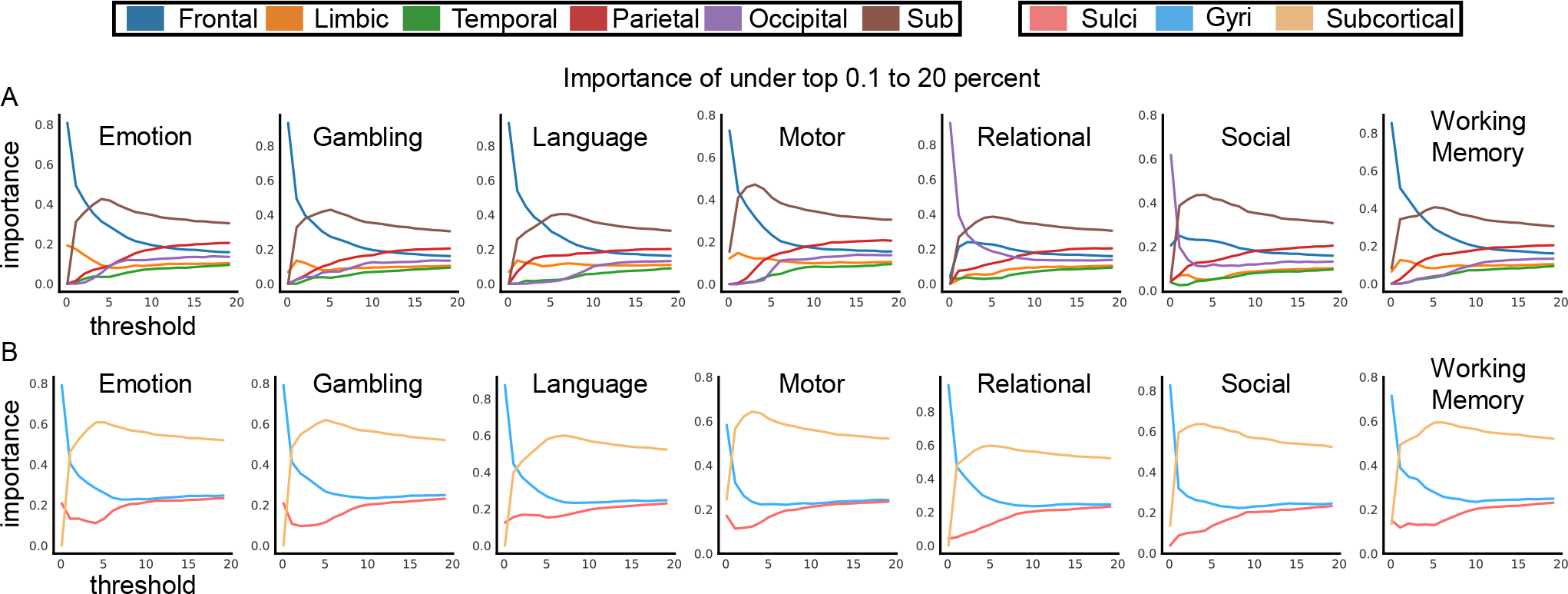
Average degree of six lobes and gyri/sulci/subcortical regions in seven tasks under different thresholds. Fig. 5A shows the average degree of six lobes. Fig. 5B shows the average degree of gyri/sulci/subcortical regions.

We then calculate the degrees of each brain region at a threshold of top 0.1% and top 5%, then select the four most important brain regions to analysis (Table I). We find that at a threshold of top 0.1%, the most important brain regions of emotion, gambling, language, motor, and working memory are all distributed in the frontal region, with medial orbitofrontal, rostral middle frontal, superior frontal, and lateral orbitofrontal being the four most important brain regions. Contrary to these five tasks, the brain regions where the important connections of relational and social located are mainly distributed in the occipital lobe at a threshold of top 0.1%, with lateral occipital, lingual and pericalcarine being the most important.

**Table I.**
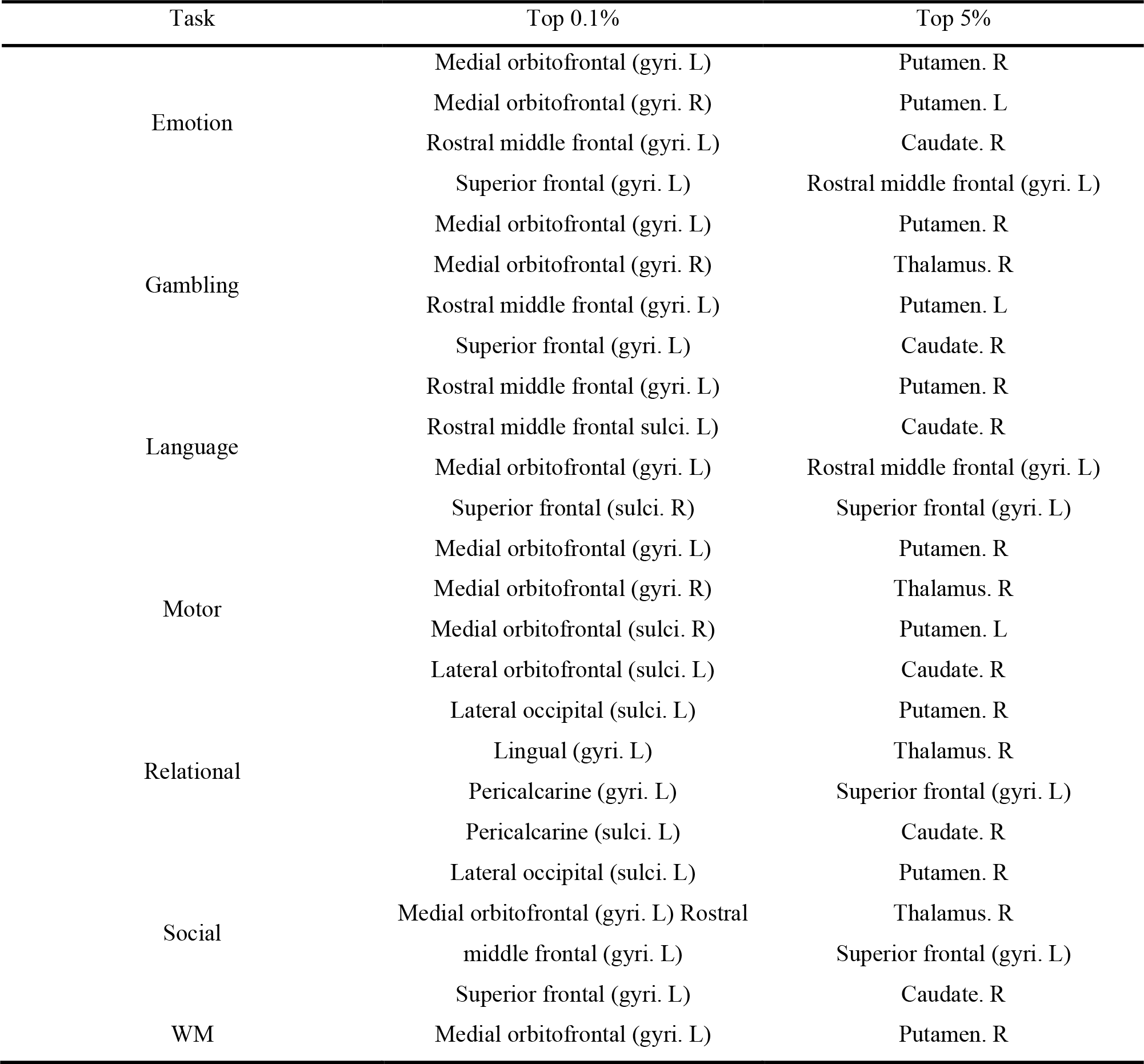
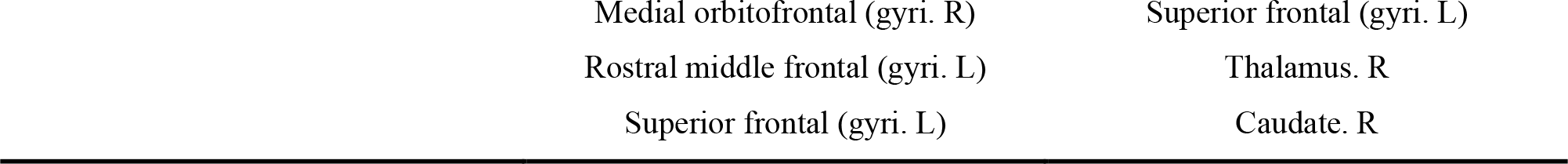
Top four brain regions in two thresholds.

As the threshold ratio increases, the brain connection pattern changes, and sub-sub connections becomes the most important. The importance of the subcortical areas is highlighted, with putamen, caudate, and thalamus widely distributed in seven task states, superior frontal and rostral middle frontal also play important roles in some tasks. These important subcortical areas are more involved in the connections between sub-sub and sub-gyri, and these connections are mostly distributed between subcortical, parietal, and frontal, indicating that gyri play an important role in regulating brain function.

## Discussion

In this study, we propose a deep learning method based on graph attention network (GAT) to explore the characteristics of structure-informed functional connectivity matrices among gyral/sulcal/subcortical regions under different tasks. To the best of our knowledge, this is the first study to combine gyri/sulci with subcortical regions to explore the brain’s functional connectivity characteristics. 999 participants from the HCP dataset, including seven task states (emotion, gambling, language, motor, relative, social, wm) and reconstructed structural connection matrices, are used as inputs to our model.

Firstly, our experimental results indicate that using GAT for classification between task and resting states can achieve an accuracy of at least 99.41%. With such high accuracy, the model can accurately identify the specificity between the task state and the resting state. Using gradient based interpretable methods, we successfully obtained important connections for each task state. We find that for the three types of connections in the cortex (gyri-gyri, gyri-sulci, sulci-sulci), gyri-gyri are the most important connections in all task states at different thresholds, gyri-sulci in the middle and sulci-sulci at the least. This is consistent with our previous findings (M. Jiang et al., 2022). Due to add subcortical regions, we find that gyri-gyri are the most important connections only at low thresholds. As the threshold increases, sub-sub becomes the most important connection, with gyri-sub in the second and sulci-sub in the third. This demonstrates the importance of the subcortical regions in different task states. In low threshold, important connections of emotion, gambling, language, motor, and working memory tasks are all distributed in the frontal, limbic and subcortical regions, while in the relationship and social tasks, they are mainly distributed in the occipital, frontal, and subcortical regions. In a higher threshold, all lobes are involved in the connections, but the three most important lobes are subcortical, parietal, and frontal. Next, we explore the common connections in different tasks and find that these common connections have stable connection patterns under different thresholds. Most of them are the connections between frontal, limbic and subcortical. In low threshold, these connections are also mainly gyri-gyri, which together with gyri-sulci and sulci-sulci constitute common connections in seven tasks.

When we use the metric of degree in graph theory to measure the important brain regions in these important connections in a low and relatively high thresholds, we find that the most important regions are not the same. In low threshold, they mainly distributed in frontal in emotion, gambling, language, motor and wm, and occipital in relational and social. In frontal, medial orbitofrontal, rostral middle frontal, superior frontal, and lateral orbitofrontal being the four most important brain regions. Orbitofrontal cortex (OFC) is widely involved in various functional activities and it has the ability of stimulate-reinforcement association learning (Rolls, 2004). OFC receives visual information from the outside and widely participates in functional activities such as decision making, emotion processing, working memory, and reward-related and punishment-related behavior (Rolls, 1996; Gallagher, McMahan, & Schoenbaum, 1999; L. Tremblay & Schultz, 1999; Bechara, Damasio, & Damasio, 2000). Recent studies have also shown that there may be functional separation between medial OFC and lateral OFC, with medial OFC being more involved in event-related fMRI (O’Doherty et al., 2003) and lateral OFC more involved in the action of suppression of previously rewarded responses(Elliott, Dolan, & Frith, 2000). This explains why the level and breadth of participation of media OFC are higher than later OFC. Middle frontal gyrus (MFG) is considered to be the intersection of dorsal and ventral attention networks, which can act as a circuit breaker to interrupt exogenic attention and redistribute attention to exogenic stimulus (Japee, Holiday, Satyshur, Mukai, & Ungerleider, 2015). At the same time, studies have shown that MFC has a regulatory effect on making mistakes. When participants lose money due to making mistakes, the hemodynamic activity response of MFC increases sharply(Taylor et al., 2006). MFG also plays an important role in working memory, serving as the center of the storage and processing part of the human brain’s working memory, as well as being responsible for the storage of spatial memory (Leung, Gore, & Goldman-Rakic, 2002). Superior frontal gyrus (SFG) plays an important role in advanced cognitive function, and previous studies have shown its significant reuse in cognition and emotional processing (Kraljević et al., 2021), gambling (Van Holst, van den Brink, Veltman, & Goudriaan, 2010), and working memory (Klingberg, 2006).

In occipital, lateral occipital, lingual and pericalcarine being the most important. Lateral occipital (LO) cortex plays an important role in object recognition, as it participates in the recognition of the shape of the object (Grill-Spector, Kourtzi, & Kanwisher, 2001). Lingual is also responsible for processing global visual feature (Mechelli, Humphreys, Mayall, Olson, & Price, 2000), and as a part of medial occipital lobe, it contribute to the cross-modal, nonvisual functions such as linguistic and verbal memory (Palejwala et al., 2021). Pericalcarine is responsible for perceiving, processing, and interpreting visual information from the retina in the visual system, and providing the brain with cognition and understanding of external visual scenarios (Silver, Ress, & Heeger, 2007). We found that gyri are mainly involved in frontal lobe, while gyri and sulci are simultaneously involved in occipital lobe, which indicates that under different tasks, gyri serve as central nodes in advanced functional regions and connect with other regions, responsible for regulating overall brain function. It is worth noting that gyri play a major role in the brain regions involved in advanced cognitive functions in the cerebral cortex, while sulci play a major role in primary functions such as visual areas. This reveals the importance of gyri in executing tasks.

When in high thresholds, subcortical regions become the most important, with putamen, caudate, and thalamus widely distributed in seven task states, superior frontal and rostral middle frontal also play important roles in some tasks. Caudate and putamen together form the basal ganglia of the brain, putamen is connected to other brain regions, forming a basal ganglia-cortical circuit network that plays an important role in motor control, learning, and memory. At the same time, it is closely connected to the limbic system and the dopamine system in the brain, controlling functions such as emotions, rewards, and addiction (Bolam, Hanley, Booth, & Bevan, 2000). Caudate plays an important role in motor control, learning, memory, and emotional regulation, and is interconnected with the prefrontal cortex in the cerebral cortex, participating in decision-making processes (Grahn, Parkinson, & Owen, 2008). Thalamus is an important nucleus located between the cerebral cortex and brainstem. It plays a crucial role in perception, motor control, emotional regulation, and consciousness (Jones, 2012).

In conclusion, this study for the first time reveals the fundamental principle of connectivity characteristics among gyri, sulci, and subcortical regions at the whole brain scale. In addition, by introducing structural connections as constraints, the functional characteristics of the brain were explored from the perspective of connectome. Our conclusion indicates that gyri play a more important role than sulci in different tasks, and the connections related to gyri are more distributed in the front and sub regional regions, revealing the importance of gyri in task regulation and its global characteristics.

## Author contributions

Conceptualization: Zifan Wang, Xi Jiang

Methodology: Zifan Wang, Paule-J Toussaint

Visualization: Zifan Wang, Xi Jiang

Funding acquisition: Xi Jiang

Project administration: Xi Jiang

Data curation: Zifan Wang

Supervision: Xi Jiang, Alan C Evans, Paule-J Toussaint

Writing – original draft: Zifan Wang

Writing – review & editing: Xi Jiang, Alan C Evans, Paule-J Toussaint

## Competing interestings

Authors declare that they have no competing interestings.

## Data and materials availability

All data are available in the main text or the supplementary materials. The HCP dataset is available in https://www.humanconnectome.org/.

